# Natural and synthetic inhibitors of a phage-encoded quorum-sensing receptor affect phage-host dynamics in mixed bacterial communities

**DOI:** 10.1101/2022.10.18.512784

**Authors:** Justin E. Silpe, Olivia P. Duddy, Bonnie L. Bassler

**Affiliations:** Department of Molecular Biology, Princeton University, Princeton, NJ, USA; Howard Hughes Medical Institute, Chevy Chase, MD, USA

## Abstract

Viruses that infect bacteria, called phages, shape the composition of bacterial communities and are important drivers of bacterial evolution. We recently showed that temperate phages, when residing in bacteria (i.e., prophages), are capable of manipulating the bacterial cell-to-cell communication process called quorum sensing (QS). QS relies on the production, release, and population-wide detection of signaling molecules called autoinducers (AI). Gram-negative bacteria commonly employ *N*-acyl homoserine lactones (HSL) as AIs that are detected by LuxR-type QS receptors. Phage ARM81ld is a prophage of the aquatic bacterium *Aeromonas* sp. ARM81, and it encodes a homolog of a bacterial LuxR, called LuxR_ARM81ld_. LuxR_ARM81ld_ detects host *Aeromonas*-produced C4-HSL, and in response, activates the phage lytic program, triggering death of its host and release of viral particles. Here, we show that phage LuxR_ARM81ld_ activity is modulated by non-cognate HSL ligands and by a synthetic small molecule inhibitor. We determine that HSLs with acyl chain lengths equal to or longer than C8 antagonize LuxR_ARM81ld_. For example, the C8-HSL AI produced by *Vibrio fischeri* that co-exists with *Aeromonads* in marine environments, binds to and inhibits LuxR_ARM81ld_, and consequently, protects the host from lysis. Co-culture of *V. fischeri* with the *Aeromonas* sp. ARM81 lysogen suppresses phage ARM81ld virion production. We propose that the cell density and species composition of the bacterial community could determine outcomes in bacterial-phage partnerships.

**SIGNIFICANCE:** Bacteria use the cell-to-cell communication process called quorum sensing to orchestrate group behaviors. Quorum sensing relies on extracellular molecules called autoinducers. Bacteria-infecting viruses (phages) can possess homologs of bacterial quorum-sensing receptors that detect autoinducers to control lysis-lysogeny transitions. We show that a phage LuxR-type quorum-sensing receptor is activated by the autoinducer produced by its host bacterium and is inhibited by non-cognate autoinducers made by bacteria that naturally co-exist with the phage’s host and by a synthetic quorum-sensing inhibitor. Our findings demonstrate that microbial community composition, mediated through quorum-sensing-communication, influences phage lysis-lysogeny transitions. These results deepen the understanding of host-phage interactions in communities and could inspire new phage-specific, quorum-sensing interventions.

## INTRODUCTION

Bacteria communicate and orchestrate collective behaviors using a process called quorum sensing (QS) (1). QS relies on the production, release, accumulation, and group-wide detection of molecules called autoinducers (AIs). Bacteria commonly live in environments containing multiple bacterial species, and thus, different blends of QS AIs can be present. Homoserine lactones (HSL) represent a common class of QS AIs produced and detected by Gram-negative bacteria. HSL AIs possess different modifications at the C3 position and they harbor variable acyl chain lengths. LuxR-type and LuxN-type QS receptors detect HSL AIs (1). Some of these QS receptors display strict specificity for a cognate HSL AI, while others are promiscuous in HSL ligand detection (2–8). For instance, *Vibrio harveyi* LuxN is exclusively activated by its partner 3OHC4-HSL ligand and non-cognate HSLs possessing longer acyl tails act as competitive antagonists (7). Such antagonism is thought to be a mechanism QS bacteria use to monitor and react to the presence of competing bacterial species. Specifically, species whose QS receptors are antagonized by non-cognate AIs repress their QS outputs when the non-cognate compounds are present, thereby avoiding leakage of QS-controlled public goods to competitors (7, 9).

Beyond QS driving interactions within and between bacterial species, we recently discovered that linear plasmid-like phages can encode LuxR-type QS receptors that detect the HSL AI produced by the bacterial host. For example, *Aeromonas* sp. ARM81 possesses a prophage, called ARM81ld, that encodes *luxR_ARM81ld_*. LuxR_ARM81ld_ binds to and is solubilized by C4-HSL, the AI made by its *Aeromonas* sp. ARM81 host (10, 11). Together with a partner XRE_ARM81ld_ DNA-binding protein, the LuxR_ARM81ld_-C4-HSL complex activates transcription of a counter-oriented gene encoding a small ORF (*smORF_ARM81ld_*) (10). Production of smORF_ARM81ld_ launches the phage ARM81ld lytic program, which causes host-cell lysis (10). Thus, monitoring its host’s QS status, via C4-HSL, allows phage ARM81ld to transition from its lysogenic to its lytic lifestyle and to disseminate at high host-cell density, presumably a condition that maximizes the probability of subsequent successful infection.

The finding that non-cognate HSLs are inhibitory to some bacterial LuxR-type and LuxN-type receptors is intriguing because it enables bacteria to take a census of and react to non-kin bacteria in the vicinity. Whether phages that possess QS receptors also detect and respond differently to non-host-produced AIs is unknown. Here, we assess the effects of non-cognate AIs on lifestyle choices made by phage ARM81ld. We demonstrate that microbial community composition, mediated through the different AIs produced, has a dramatic influence on phage ARM81ld lysis-lysogeny transitions. These results have potentially far-reaching implications for how we understand host-phage interactions in complex communities and could lead to the development of new classes of QS-targeted interventions that are phage- rather than bacteria-specific.

## RESULTS

*Aeromonads* are known to exist in mixed microbial consortia with other QS bacteria, particularly marine *Vibrios*. *Vibrio fischeri* is one such well-studied QS bacterium. It produces 3OC6-HSL and C8-HSL (12, 13), two AIs that have longer acyl tails than the C4-HSL AI to which phage ARM81ld responds. We verified that C4-HSL is the product of the *Aeromonas* sp. ARM81 AhyI AI synthase using an established bioassay (Figure S1A) (10, 14). To explore the effects of signaling molecules that *Aeromonas* sp. ARM81 encounters in communities but that it itself does not produce on phage ARM81ld activity, we constructed an *Aeromonas* sp. ARM81 lysogen that was incapable of producing C4-HSL to eliminate any phage activity that occurs in response to the endogenously-produced AI. We used this strain (designated Δ*ahyI Aeromonas* sp. ARM81) in all of our assays. We introduced anhydrotetracyline (aTc)-inducible *xre_ARM81ld_-luxR_ARM81ld_* on a plasmid into Δ*ahyI Aeromonas* sp. ARM81. We induced production of XRE_ARM81ld_-LuxR_ARM81ld_ and administered cell-free culture fluids collected from wild-type (WT) *V. fischeri*. As a control, we administered cell-free fluids from WT *E. coli*, which does not produce HSL AIs. Important for our strategy is that we used a concentration of the aTc inducer sufficient to drive an intermediate-level of phage-directed host-cell lysis in the absence of exogenous ligand, thus enabling us to detect increased or decreased cell death (Figure 1A). Strikingly, culture fluids from WT *V. fischeri* completely suppressed cell lysis (Figure 1A). By contrast, culture fluids from WT *E. coli* did not affect cell lysis relative to medium alone (Figure 1A).

**Figure 1.**
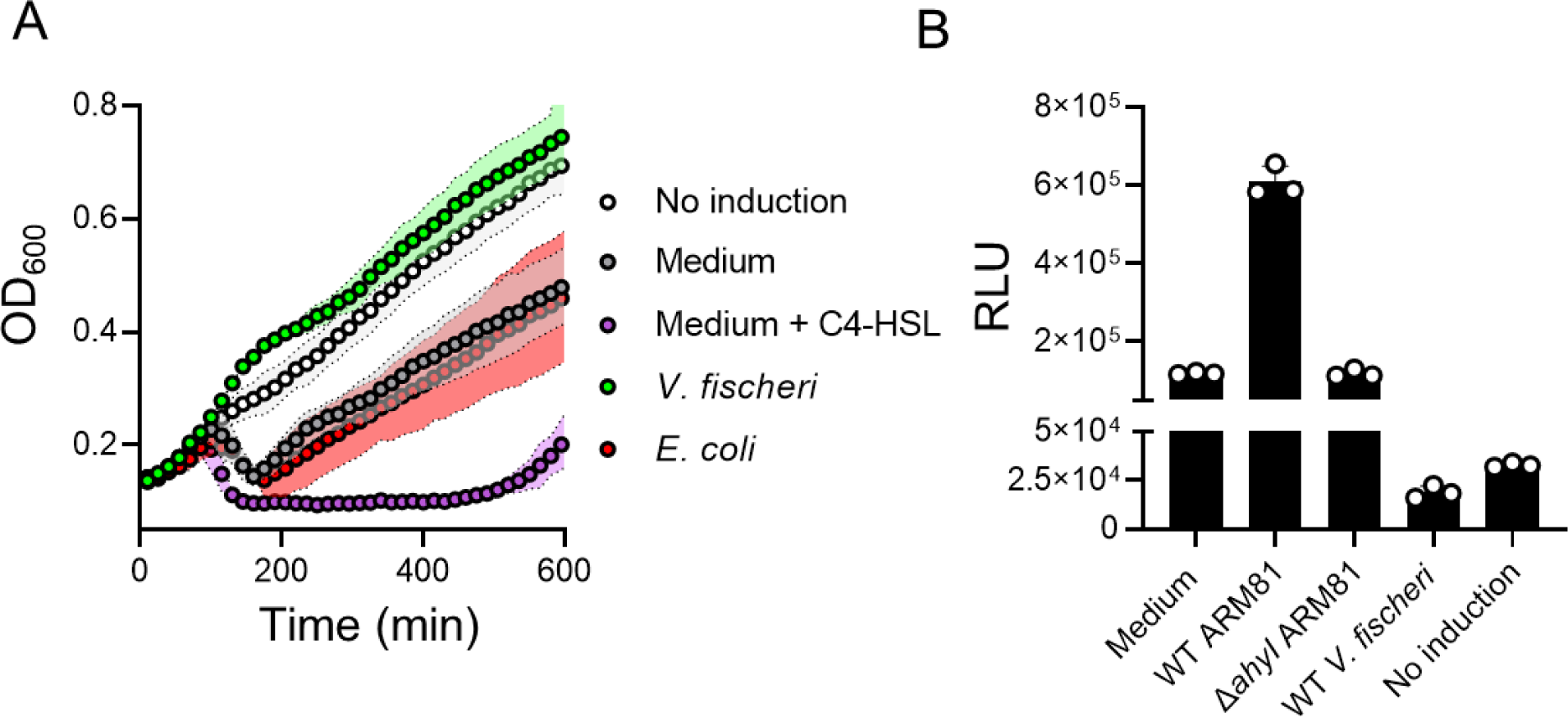
*V. fischeri* culture fluids inhibit XRE_ARM81ld_-LuxR_ARM81ld_ transcriptional activity. (a) Growth of Δ*ahyI Aeromonas* sp. ARM81 carrying aTc-inducible *xre_ARM81ld_-luxR_ARM81ld_* in medium lacking aTc (white; No induction), medium containing 0.1 ng mL^−1^ aTc supplemented with medium alone (gray), medium and 20 μM C4-HSL (purple), culture fluids from WT *V. fischeri* (3OC6-HSL^+^ and C8-HSL^+^; green), or culture fluids from *E. coli* (a non-HSL producer; red). (b) P*smORF_ARM81l_d*-*lux* expression from *E. coli* carrying aTc-inducible *xre_ARM81ld_-luxR_ARM81ld_* grown in medium containing 50 ng mL^−1^ aTc supplemented with medium alone, culture fluids from WT *Aeromonas* sp. ARM81 (C4-HSL^+^), culture fluids from Δ*ahyI Aeromonas* sp. ARM81 (C4-HSL^−^), culture fluids from WT *V. fischeri* (3OC6-HSL^+^ and C8-HSL^+^), or in medium lacking aTc (No induction). RLU denotes relative light units. Data are represented as means ± std with *n*=3 biological replicates.

Given that XRE_ARM81ld_-LuxR_ARM81ld_-mediated transcription of *smORF_ARM81ld_* drives host-cell lysis by phage ARM81ld (10), we hypothesized that the inhibitory effect of *V. fischeri* culture fluids occurred through suppression of XRE_ARM81ld_-LuxR_ARM81ld_ transcriptional activity. To explore this possibility, we used recombinant *E. coli* harboring a P*smORF_ARM81ld_-lux* transcriptional reporter and aTc-inducible *xre_ARM81ld_-luxR_ARM81ld_,* thus excluding all other *Aeromonas* sp. ARM81 host-and phage components from the system. Consistent with our understanding that C4-HSL activates the XRE_ARM81ld_-LuxR_ARM81ld_ pathway and occurs at concentrations relevant to that produced by *Aeromonas* sp. ARM81 in nature, administration of culture fluids from WT *Aeromonas* sp. ARM81 increased P*smORF_ARM81ld_-lux* light production 5-fold over medium alone, whereas culture fluids from Δ*ahyI Aeromonas* sp. ARM81 had no effect (Figure 1B). Importantly, culture fluids from WT *V. fischeri* inhibited light production 6-fold, indeed to levels below that when expression of *xre_ARM81ld_-luxR_ARM81ld_* was not induced (Figure 1B). Thus, *V. fischeri* culture fluids harbor a factor(s) that prevents phage-mediated cell lysis by inhibiting the phage-encoded XRE_ARM81ld_-LuxR_ARM81ld_ pathway.

As noted, *V. fischeri* makes two HSL AIs, 3OC6-HSL and C8-HSL. To test whether the inhibition of *Aeromonas* sp. ARM81 lysis shown in Figure 1A is due to one or both of these AIs, we administered cell-free culture fluids harvested from Δ*luxI V. fischeri*, which makes no 3OC6-HSL, Δ*ainS V. fischeri* which makes no C8-HSL, and Δ*luxI* Δ*ainS V. fischeri* which makes neither AI to the *Aeromonas* sp. ARM81 lysogen (15–17). Identical to the case of WT *V. fischeri* culture fluids, addition of culture fluids from Δ*luxI V. fischeri* inhibited *Aeromonas* sp. ARM81 lysis. By contrast, culture fluids from Δ*ainS* or Δ*luxI* Δ*ainS V. fischeri* only drove basal-level lysis of *Aeromonas* sp. ARM81, i.e., to the same level as when medium alone was added (Figure 2A). Consistent with this result, WT and Δ*luxI* culture fluids decreased P*smORF_ARM81ld_-lux* output 10-fold while Δ*ainS* and Δ*luxI* Δ*ainS* culture fluids had a <2-fold effect (Figure 2B). These findings suggest that the *V. fischeri* AIs, primarily C8-HSL, antagonize LuxR_ARM81ld_. Indeed, administration of synthetic C8-HSL to the *Aeromonas* sp. ARM81 lysogen inhibited cell lysis and decreased reporter output 8-fold (Figures 2C and 2D, respectively). By comparison, synthetic 3OC6-HSL had no effect on lysis and a modest activating effect (2.5-fold) on P*smORF_ARM81ld_* expression (Figure 2C and 2D, respectively). Maximum cell lysis and maximum activation of the reporter by C4-HSL are shown as controls (Figure 2C and 2D, respectively). Likely, C8-HSL is a more potent antagonist than 3OC6-HSL is an agonist of LuxR_ARM81ld_. Thus, C8-HSL is the *V. fischeri* AI that protects the *Aeromonas* sp. ARM81 lysogen from undergoing phage-mediated lysis.

**Figure 2.**
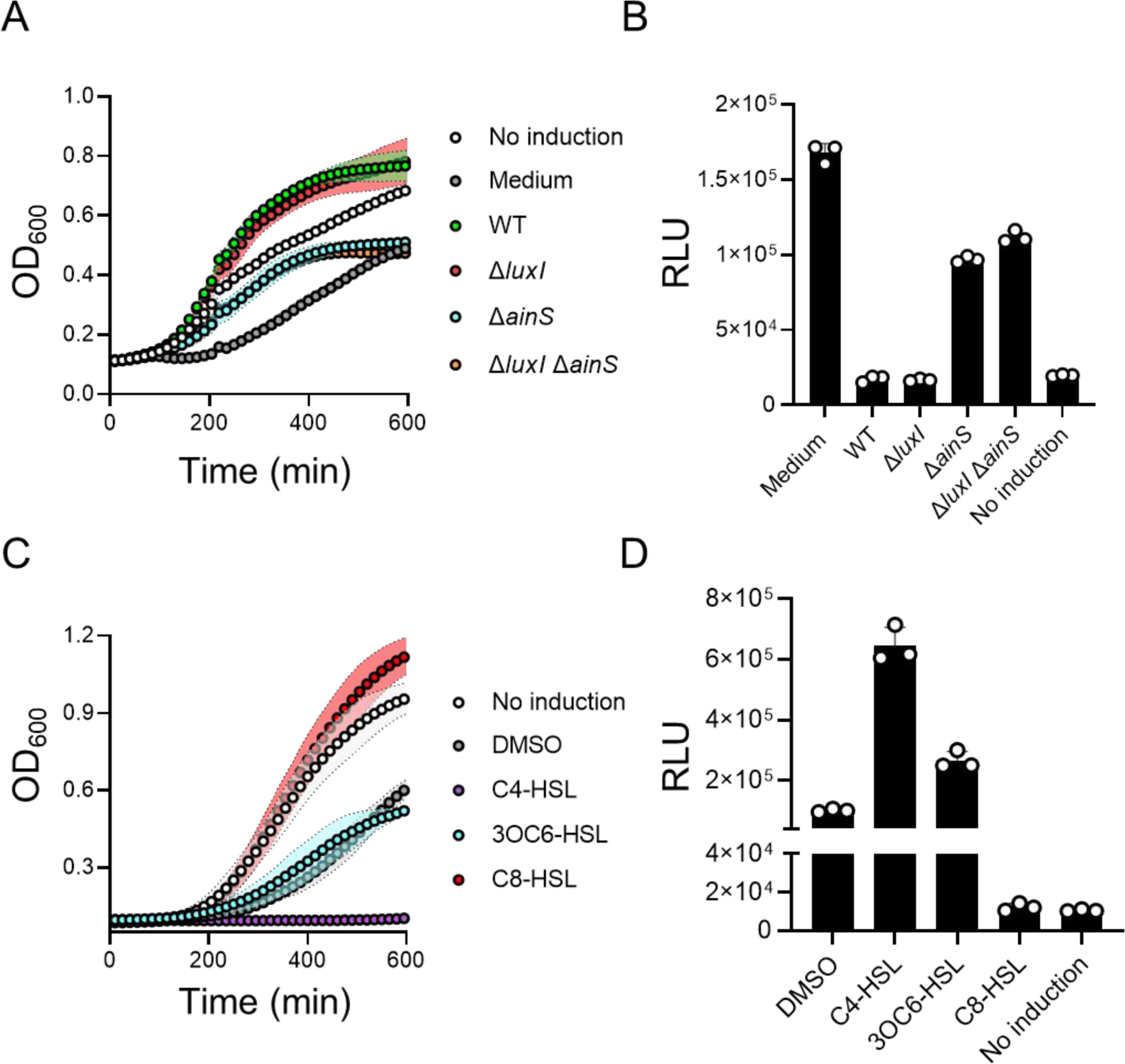
The *V. fischeri* C8-HSL AI antagonizes LuxR_ARM81ld_. (a) Growth of Δ*ahyI Aeromonas* sp. ARM81 carrying aTc-inducible *xre_ARM81ld_-luxR_ARM81ld_* in medium lacking aTc (white; No induction), medium containing aTc supplemented with medium alone (gray), culture fluids from WT (3OC6-HSL^+^ and C8-HSL^+^; green), Δ*luxI* (3OC6-HSL^−^ and C8-HSL^+^; red), Δ*ainS* (3OC6-HSL^+^ and C8-HSL^−^; cyan), or Δ*luxI* Δ*ainS* (3OC6-HSL^−^ and C8-HSL^−^; orange) *V. fischeri*. (b) P*smORF_ARM81ld_*-*lux* expression from *E. coli* carrying aTc-inducible *xre_ARM81ld_-luxR_ARM81ld_* grown in medium containing aTc supplemented with medium alone, culture fluids from WT (3OC6-HSL^+^ and C8-HSL^+^) *V. fischeri*, Δ*luxI* (3OC6-HSL^−^ and C8-HSL^+^) *V. fischeri*, Δ*ainS* (3OC6-HSL^+^ and C8-HSL^−^) *V. fischeri*, Δ*luxI* Δ*ainS* (3OC6-HSL^−^ and C8-HSL^−^) *V. fischeri*, or in medium lacking aTc (No induction). (c) Growth of Δ*ahyI Aeromonas* sp. ARM81 carrying aTc-inducible *xre_ARM81ld_-luxR_ARM81ld_* in medium lacking aTc (white; No induction), medium containing aTc supplemented with DMSO (gray), C4-HSL (purple), 3OC6-HSL (cyan) or C8-HSL (red). All HSLs were supplied at 20 μM. (d) P*smORF_ARM81ld_*-*lux* expression from *E. coli* carrying aTc-inducible *xre_ARM81ld_-luxR_ARM81ld_* grown in medium containing aTc supplemented with DMSO, C4-HSL, 3OC6-HSL, C8-HSL, or in medium lacking aTc (No induction). HSL concentrations as in (c). Data are represented as means ± std with *n*=3 biological replicates. RLU as in Figure 1b (b, d). aTc; 0.1 ng mL^−1^ (a, c), 50 ng mL^−1^ (b) and 25 ng mL^−1^ (d).

Despite our finding that C4-HSL promotes and C8-HSL prevents XRE_ARM81ld_-LuxR_ARM81ld_-driven host-cell lysis, both HSLs solubilize LuxR_ARM81ld_ (Figure S1B) (11). We thus wondered what features of HSL ligands distinguish inhibition from activation of LuxR_ARM81ld_. To probe this question, we administered a panel of synthetic HSLs to the *E. coli* P*smORF_ARM81ld_-lux* reporter (Figure 3A and 3B). Light output increased in the presence of C4-HSL, 3OC4-HSL, and 3OC6-HSL (Figure 3B). C6-HSL had no effect (Figure 3B). Conversely, HSLs with chain lengths of C8 or longer reduced P*smORF_ARM81d_* expression 5-7-fold relative to the basal activity generated by the presence of XRE_ARM81ld_ and LuxR_ARM81ld_ (Figure 3B). We also assayed the compound meta-bromo-thiolactone (mBTL, Figure 3A), a synthetic inhibitor of LuxR-driven QS (18). Similar to the longer acyl chain HSL AIs, mBTL inhibited P*smORF_ARM81ld_-lux* activity 6.5-fold (Figure 3B). Finally, simultaneous administration of the C4-HSL agonist and the C8-HSL antagonist revealed that LuxR_ARM81ld_ is highly sensitive to and prefers C4-HSL, but C8-HSL can compete for binding when provided at 60-250-fold higher concentrations (Figure 3C). This result is consistent with the finding that C4-HSL solubilizes LuxR_ARM81ld_ more effectively than C8-HSL (Figure S1B) (11).

**Figure 3.**
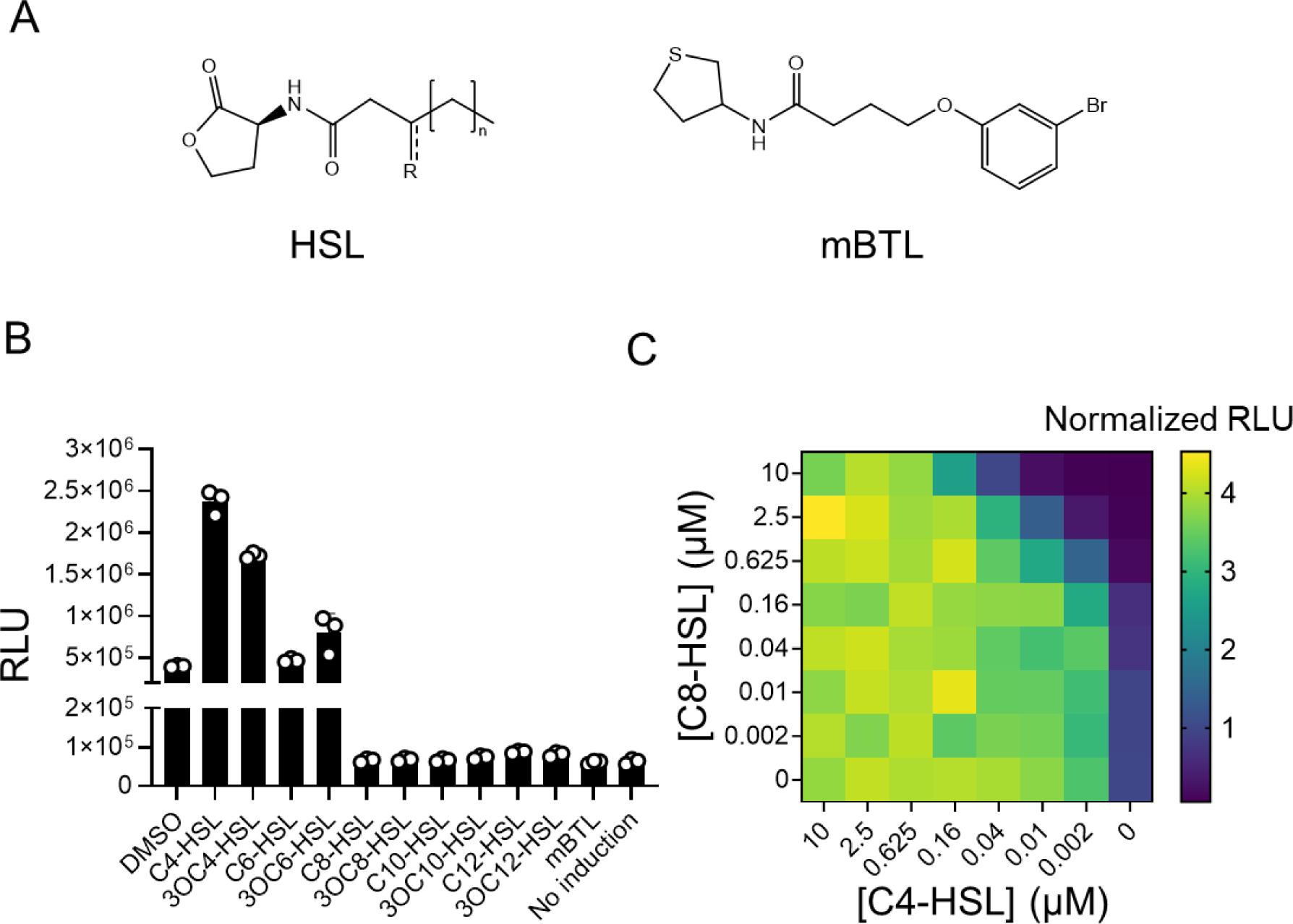
Non-cognate HSL AIs with chain lengths of C8 or longer, and the synthetic compound mBTL, inhibit LuxR_ARM81ld_ activity. (a) General structure of an HSL AI (R = O or H; n = 0, 2, 4, 6, or 8) and the structure of the synthetic compound mBTL. (b) P*smORF_ARM81ld_*-*lux* expression from *E. coli* carrying aTc-inducible *xre_ARM81ld_-luxR_ARM81ld_* grown in medium containing aTc supplemented with DMSO or the indicated compounds or in medium lacking aTc (No induction). HSL concentrations as in Figure 2c. (c) P*smORF_ARM81ld_*-*lux* expression from *E. coli* carrying aTc-inducible *xre_ARM81ld_-luxR_ARM81ld_* grown in medium containing aTc and the indicated concentrations of C4-HSL and C8-HSL. Data are shown as a heatmap. Normalized RLU refers to the RLU of each sample relative to the RLU of the sample administered DMSO only, which was set to 1.0. Data are represented as means ± std with *n*=3 biological replicates (b) or as means with *n*=2 biological replicates (c). RLU as in Figure 1b (b, c). aTc; 25 ng mL^−1^ (b, c).

Our above results imply that in mixed-species communities, whether the *Aeromonas* sp. ARM81 lysogen is killed by or protected from its prophage partner could depend on whether other species in the vicinal community are QS-proficient bacteria or not, and if the former, on what particular HSLs they produce. To garner evidence for this notion, we grew the Δ*ahyI Aeromonas* sp. ARM81 lysogen harboring inducible *xre_ARM81ld_-luxR_ARM81ld_*, alone or in combination with either WT, Δ*luxI*, Δ*ainS*, or Δ*luxI* Δ*ainS V. fischeri*. Quantitation of the ARM81ld phage-to-host ratio revealed that the viral load was approximately 4-fold lower when Δ*ahyI Aeromonas* sp. ARM81 was grown in co-culture with *V. fischeri* that produce C8-HSL (WT and Δ*luxI V. fischeri*) than when Δ*ahyI Aeromonas* sp. ARM81 was grown in mono-culture or in co-culture with *V. fischeri* strains that lacked the ability to produce C8-HSL (Δ*ainS* or Δ*luxI* Δ*ainS V. fischeri*) (Figure 4, black bars). A similar trend but, not surprisingly, with a reduced effect occurred when the WT *Aeromonas* sp. ARM81 lysogen that produces endogenous C4-HSL was used (Figure 4, white bars). Together, these results indicate that, under the conditions tested, the presence of *V. fischeri* suppresses induction of phage ARM81ld and diminishes release of phage particles, including from the C4-HSL producing (WT) *Aeromonas* sp. ARM81 lysogen. The inhibitory effect relies on *V. fischeri* production of C8-HSL and operates by C8-HSL antagonism of the phage-encoded QS receptor in the neighboring *Aeromonas* sp. ARM81 lysogen.

**Figure 4.**
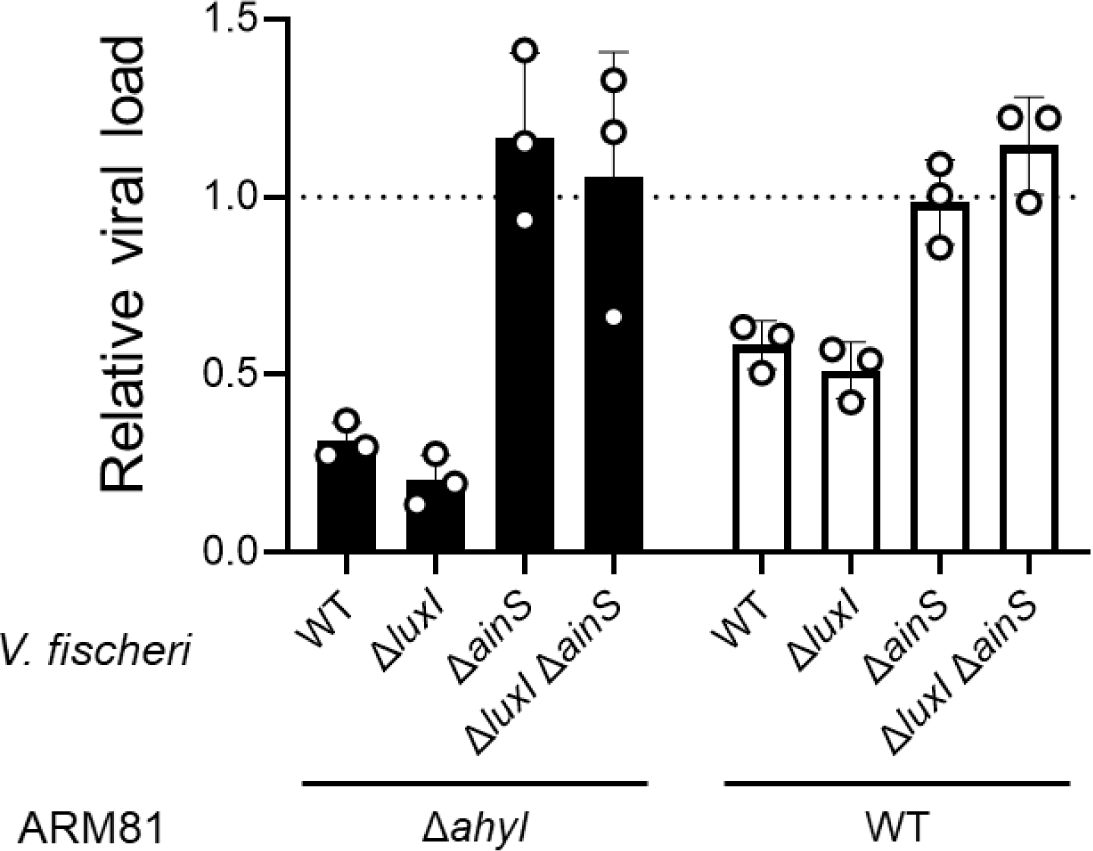
*V. fischeri* that produces C8-HSL prevents phage ARM81ld-driven virion production in co-culture with the Δ*ahyI* and WT ARM81 lysogens. Detection of phage ARM81ld obtained from cultures of Δ*ahyI Aeromonas* sp. ARM81 (black bars) or WT *Aeromonas* sp. ARM81 (white bars) carrying aTc-inducible *xre_ARM81ld_-luxR_ARM81ld_* that were grown in co-culture with the indicated *V. fischeri* strains. Relative viral load is the amount of phage ARM81ld DNA in a sample compared to the amount of *Aeromonas* sp. ARM81 host DNA. Data are represented as means ± std with *n*=3 biological replicates and *n*=3 technical replicates. aTc; 0.1 ng mL^−1^.

## DISCUSSION

Here, we show that the outcome of the phage ARM81ld lysis-lysogeny transition can be altered by other bacterial species in the community that engage in QS and produce non-cognate HSL AIs. Our findings suggest that phage ARM81ld monitors its host’s QS status and also the cell density and species composition of the vicinal community. The information it garners exists in the form of QS chemical cues, and it integrates that information into its lysis-lysogeny decision-making mechanism. We propose that, in communities in which multiple bacterial species and phages co-exist, detection of a variety of HSL AIs could benefit the phage or the host and which entity receives the benefit likely depends on the particular circumstances. First, regarding a possible benefit to the phage: Antagonism of LuxR_ARM81ld_ by non-cognate HSL AIs could prevent premature launch of the phage ARM81ld lytic cascade, and release of viral particles under conditions where *Aeromonad*s make up only a minority of a mixed-species community. Because the phage ARM81ld host range is likely limited to *Aeromonad*s, this mechanism could prevent phage ARM81ld from launching its lytic cycle when the likelihood of released virions encountering suitable bacteria to infect is low. Alternatively, regarding a possible benefit to the *Aeromonas* host: The production of non-cognate AIs by other members of the vicinal bacterial community could suppress QS-mediated induction of the *Aeromonas* sp. ARM81 lysogen, curb release of phage ARM81ld virions, and thereby protect existing *Aeromonads* harboring prophages from killing as well as protect neighboring susceptible *Aeromonads* from infection.

Beyond exploring the effects of non-cognate AIs on phages in bacterial communities, we demonstrated that the synthetic mBTL compound antagonizes XRE_ARM81ld_-LuxR_ARM81ld_ transcriptional activity, and in doing so, prevents lysis of *Aeromonas* sp. ARM81. This finding suggests that synthetic molecules designed against bacterial QS systems may have significant and unintended consequences on prophages and other mobile genetic elements that may not be present in all isolates. As we continue to uncover diverse and unexpected roles phages play in biology, the ability to develop small molecules to manipulate phage-specific rather than bacteria-specific activities may be useful. Discovering and characterizing new phage regulatory systems, like that of phage ARM81ld, could be an important step for consideration in advancing this goal.

## MATERIALS & METHODS

### Bacterial strains and growth conditions

*E. coli* strains were grown with aeration in Luria-Bertani (LB-Miller, BD-Difco) broth. *Aeromonas* sp. ARM81 and *V. fischeri* strains were grown in LB with 3% NaCl. All strains were grown at 30° C. Strains used in the study are listed in Table S1. Unless otherwise noted, the following antibiotics and concentrations were used: 100 μg mL^−1^ ampicillin (Amp, Sigma), 50 μg mL^−1^ kanamycin (Kan, GoldBio), and 5 μg mL^−1^ chloramphenicol (Cm, Sigma). Inducers were used as follows: *E. coli*: 200 μM isopropyl beta-D-1-thiogalactopyranoside (IPTG, GoldBio), 0.1% L-arabinose (Sigma), and 50 ng mL^−1^ or 25 ng mL^−1^ anhydrotetracycline (aTc, Clontech) and *Aeromonas* sp. ARM81: 0.1 ng mL^−1^ aTc. HSL AIs were supplied at a final concentration of 20 μM, unless otherwise indicated.

### Cloning techniques

All primers and dsDNA (gene blocks) used for plasmid construction and qPCR, listed in Table S2, were obtained from Integrated DNA Technologies. Gibson assembly, and traditional cloning methods were employed for all constructions. PCR with iProof was used to generate insert and backbone DNA. Gibson assembly relied on the HiFi DNA assembly mix (NEB). All enzymes used in cloning were obtained from NEB. Plasmids used in this study are listed in Table S3. Transfer of plasmids into *Aeromonas* sp. ARM81 was carried out by conjugation followed by selective plating on LB supplemented with Kan and Cm.

### Lysis and reporter assays

For Δ*ahyI Aeromonas* sp. ARM81 growth and lysis assays, overnight cultures were back-diluted 1:50 with fresh medium and appropriate antibiotics before being dispensed into 96-well plates (Corning Costar 3904). Cultures were grown in the plates for 60 min prior to administration of aTc, cell-free culture fluids, HSLs, or mBTL. *E. coli* reporter assays were carried out as above with the following modifications: Overnight cultures were back-diluted 1:100 with fresh medium and appropriate antibiotics, dispensed into 96-well plates, and immediately supplied aTc, cell-free culture fluids, or HSLs. In all cases, cell-free culture fluids were administered at 30% (w/v), and plate wells that did not receive treatment received equal volumes of growth medium or DMSO, as specified. To make cell-free culture fluids, overnight cultures of *V. fischeri*, *Aeromonas* sp. ARM81, and *E. coli* strains were grown in LB + 3% NaCl and cells were removed by centrifugation. The clarified supernatants were collected and filtered through 0.22 μM filters (Corning SpinX). A BioTek Synergy Neo2 Multi-Mode reader was used to measure OD_600_ and bioluminescence. Relative light units (RLU) were calculated by dividing the bioluminescence readings by the OD_600_ reading at that time.

### Total protein and in-gel HALO detection to assess phage LuxR_ARM81ld_ solubility

Overnight cultures of *E. coli* T7Express lysY/I^q^ carrying the plasmid with the LuxR_ARM81ld_-HALO-HIS fusion were diluted 1:200 in 15 mL medium and grown at 37°C to OD_600_ ~ 0.5. 200 μM IPTG was added to each culture before it was divided into 3 equal volumes, and the aliquots received 75 μM C4-HSL, 75 μM C8-HSL, or an equivalent volume of DMSO. The cultures were returned to growth at 37 °C for an additional 3 h prior to cell collection by centrifugation. Pellets were stored at −80 °C prior to processing. Cell pellets were resuspended in a lysis buffer containing BugBuster, benzonase, and 1 μM HALO-Alexa_660_ (excitation/emission: 663/690 nm) and incubated at room temperature for 15 min. The resulting whole cell lysates were loaded onto 4-20% SDS-PAGE stain-free gels, which were imaged using an ImageQuant LAS 4000 imager under the Cy5 setting for HALO-Alexa_660_ before being exposed to UV-light for 7 min and re-imaged under the EtBr setting for total protein. Exposure times never exceeded 30 sec.

### qPCR measurement of relative phage ARM81ld viral load from co-cultures

Triplicate colonies of WT and Δ*ahyI Aeromonas* sp. ARM81 and *V. fischeri* strains were each resuspended in 1 mL fresh growth medium and incubated at 30 °C until the cultures reached OD_600_ ~ 0.5. Cultures were back-diluted 1:100 into fresh growth medium and combined at a 1:5 ratio *Aeromonas* sp. ARM81:*V. fischeri*. The *Aeromonas* sp. ARM81 mono-culture control was prepared in parallel by dilution of the *Aeromonas* sp. ARM81 culture 1:5 in growth medium. The mono- and co-cultures were dispensed into a 96-well plate and incubated at 30 °C with shaking for ~10 h, at which point 10 uL aliquots were collected, heated to 95 °C for 10 min, and diluted 1:1000 in water. SYBR Green mix (Quanta) and the Applied Biosystems QuantStudio 6 Flex Real-Time PCR detection system (Thermo) were used for real-time PCR. Data were processed and analyzed (Pfaffl method) by comparing the relative amplification within samples from reactions using an ARM81ld phage-specific primer pair (targeting *cI_ARM81ld_*, Table S3) to that from reactions using an *Aeromonas* sp. ARM81 host-specific primer pair (targeting *rpoB*, Table S3). The relative phage ARM81ld viral load was determined by dividing the ARM81 phage-to-host amplification ratio from each co-culture condition by that of the *Aeromonas* sp. ARM81 mono-culture.

### Quantitation and statistical analyses

Software used to acquire and analyze data generated in this study consisted of: GraphPad Prism9 for analysis of growth- and reporter-based experiments; Gen5 for collection of growth- and reporter-based data; SnapGene v6 for primer design; QuantStudio for qPCR quantitation; and FIJI for image analyses. Data are presented as the means ± std. The numbers of independent biological replicates for each experiment are indicated in the figure legends.

## Acknowledgments

We thank all members of the Bassler lab for insightful discussions. This work was supported by the Howard Hughes Medical Institute, National Science Foundation grant MCB-2043238, and the National Institutes of Health grant R37GM065859 to B.L.B. J.E.S. is a Howard Hughes Medical Institute Fellow of the Jane Coffin Childs Memorial Fund for Medical Research. O.P.D. was supported by the NIGMS T32GM007388 grant. The content is solely the responsibility of the authors and does not necessarily represent the official views of the funders.

## Data availability interests

All growth data, reporter data, and unprocessed gels presented in each panel of this study are provided in Table S4 and available on Zenodo (doi: 10.5281/zenodo.7209272).

## Author contributions

J.E.S., O.P.D., and B.L.B. conceptualized the project. J.E.S. constructed strains. J.E.S. and O.P.D. performed experiments. J.E.S., O.P.D., and B.L.B. analyzed data. J.E.S., O.P.D., and B.L.B. designed experiments. J.E.S., O.P.D., and B.L.B. wrote the paper.

## SUPPLEMENTARY FIGURE

**Figure S1.**
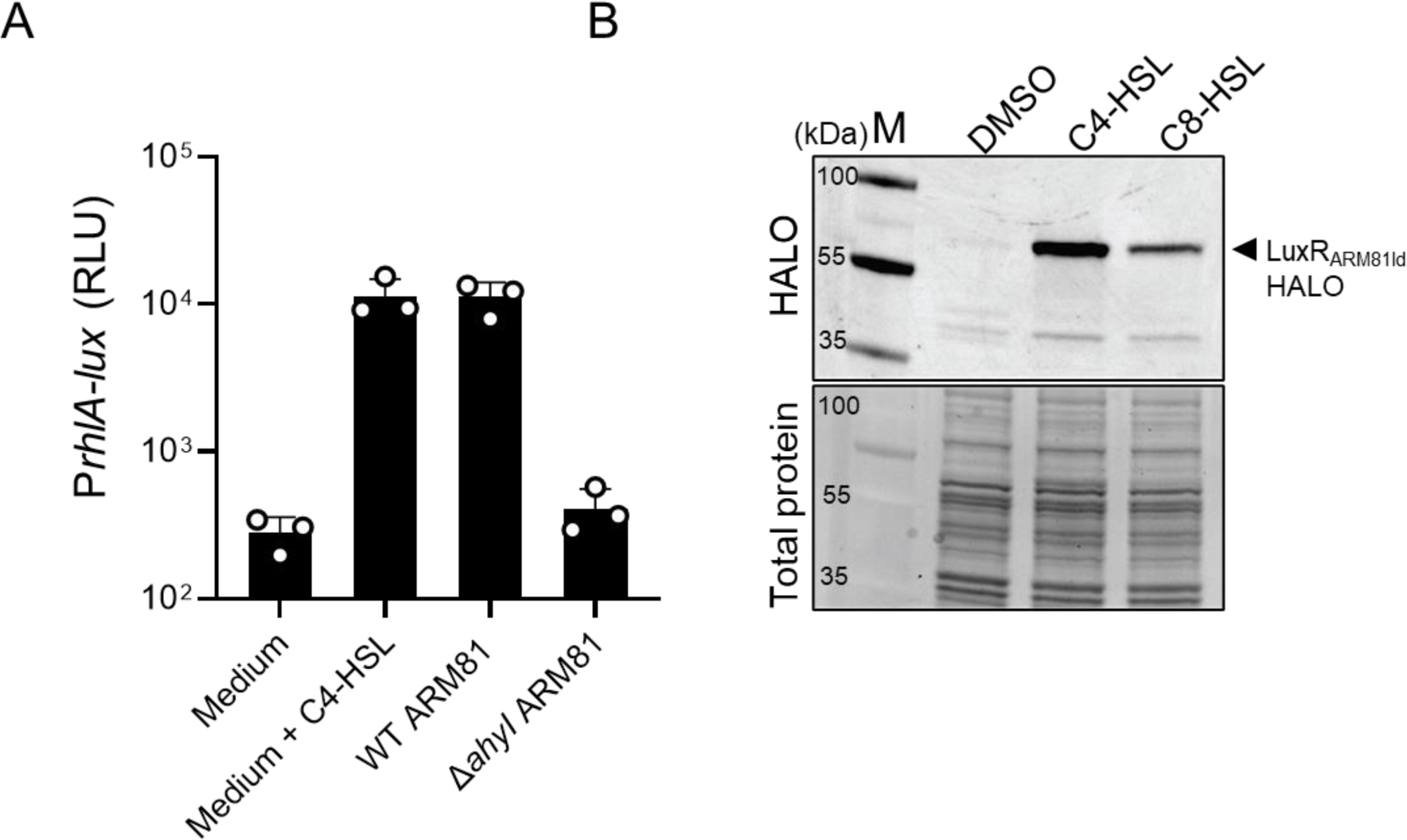
The ARM81 *ahyI* gene encodes the C4-HSL-producing synthase AhyI and both C4-HSL and C8-HSL solubilize LuxR_ARM81ld_. (A) P*rhlA*-*lux* expression from *E. coli* grown in medium supplemented with C4-HSL, culture fluids from WT *Aeromonas* sp. ARM81 (C4-HSL^+^), or culture fluids from Δ*ahyI Aeromonas* sp. ARM81 (C4-HSL^−^). The *E. coli* strain carries a plasmid with P*rhlA*-*lux* and a second plasmid with arabinose-inducible pBAD-*rhlR*. The *rhlA* promoter and *rhlR* gene both come from *P. aeruginosa.* In this system, RhlR only activates P*rhlA*-*lux* expression when C4-HSL is supplied exogenously. All media contained 0.1% arabinose. (B) Western blot (top) and total protein (bottom) showing LuxR_ARM81ld_-HALO in the soluble fractions of *E. coli* supplied with 75 μM of the indicated HSL or an equivalent volume of DMSO. Detection of the HALO-reactive fluorophore requires that the HALO tag be properly folded. Thus, LuxR_ARM81ld_-HALO band intensity is a measure of folded LuxR_ARM81ld_ in the soluble fraction of each sample. M denotes molecular weight marker (representative bands are labeled). kDa is kilodalton.

